# Energetic and functional development of soil food webs along secondary succession

**DOI:** 10.64898/2026.11.05.762856

**Authors:** Ajuan Zhang, Chengwei Tu, Yuanfei Wang, Jingyi Li, Zheng Zhou, Ruyi Luo, Karin Hohberg, Zhe Chen, Bingxue Zhu, Xueyong Pang, Nico Eisenhauer, Anton M. Potapov

## Abstract

1. As ecosystems develop, aboveground biomass and belowground organic matter accumulate, while energy turnover declines. This trend is largely supported by vegetation studies, but remains unexplored for soil food webs that channel the majority of energy in terrestrial ecosystems.
2. Here, we tested whether successional theory applies to soil food webs across size-based compartments, ranging from microfauna (nematodes) to macrofauna (soil arthropods and earthworms), along a secondary succession in southwestern China, using an energy-flux approach. To assess whether forest restoration by planting follows natural succession, we also compared spruce and larch plantations with secondary birch forest of the same successional stage.
3. Supporting expectations of reduced energy turnover, the results showed an increasing animal biomass-to-energy flux ratio and a functional shift from herbivory-dominated (green) to detritivory-dominated (brown) energy channelling along succession. Soil food web compartments showed contrasting trends: macro-food webs shifted towards bacterial energy channels with decreasing predation, while micro-food webs maintained similar bacterial/fungal energy flux ratios and tended towards increasing predation. The larch plantation supported lower animal biomass than the secondary birch forest, whereas the spruce plantation did not differ. We further found that energy flux correlated positively with microbial biomass and soil animal diversity, but negatively with litter biomass during succession.
4. Overall, we expand successional theory by revealing the transformations in energy use and pathways within soil food webs along secondary succession, thereby advancing understanding of ecosystem functioning and guiding ecosystem restoration. Our results further suggest that tree plantations do not necessarily replicate the belowground development of natural succession, which should be considered in ecosystem restoration.

## INTRODUCTION

Unravelling the mechanistic links between ecological succession and energetics in natural communities has been a recurring focus in ecology for much of the past century (Odum 1969; Poorter *et al*. 2023). Odum’s theory of ecosystem development predicts that, as succession proceeds, both aboveground biomass and belowground organic matter accumulate, the biomass supported per unit of energy flow increases, and food chains shift from grazing- to detritus-dominated pathways (Odum 1969). Despite the foundational role of this concept in succession theory, most supporting evidence is derived from studies of vegetation, even though approximately 90% of energy (net primary productivity) is directed into soil food webs (Cebrian 1999). Yet the assumption of energetic development of consumer communities in soil has been largely untested due to the intricate nature and vast biodiversity of soil organisms (Anthony *et al*. 2023), along with the challenges in comprehensive soil food web quantification (Jochum & Eisenhauer 2022).

During succession, changes in structure and increasing complexity of vegetation alter basal resources and modify environmental conditions, potentially leading to shifts in belowground communities and trophic interactions (Hooper *et al*. 2000). Although many studies have examined how belowground biodiversity and community composition respond to succession (Frouz *et al*. 2001, 2008), the organization of soil food webs and the energy fluxes through them remain poorly understood, despite their central role in linking community changes to ecosystem functioning (Barnes *et al*. 2014, 2018). Total energy flux, estimated as the sum of energy flowing through the soil food web, indicates the biological activity of soil organisms, whereas the total biomass of all measured soil animal groups—ranging from microfauna to macrofauna—represents the energy stored in the system and the resources available to consumers (Buzhdygan *et al*. 2020). Given that energy flux and energy storage (biomass) are not necessarily correlated (Barnes *et al*. 2014; Buzhdygan *et al*. 2020), it is essential to consider both measures to develop a more comprehensive understanding of successional changes. The biomass-to-energy flux ratio (B/E) quantifies the standing biomass maintained per unit energy throughput and thus approximates turnover time; Odum (1969) predicted that it increases as ecosystems mature. A higher B/E indicates slower turnover and greater retention of energy in biomass, which has been interpreted as a more efficient use of energy, but may also reflect lower metabolic activity or productivity (Buzhdygan *et al*. 2020; Odum 1969). Although energy fluxes are closely linked to biodiversity (Barnes *et al*. 2014, 2018), it remains unclear whether, along succession, energy fluxes in soil food webs track changes in basal resources, in biodiversity, or both.

Energy fluxes can serve as proxies for the functional roles of trophic interactions on ecosystem functioning – such as feeding on living plants (herbivory), dead organic matter (detritivory), microorganisms (microbivory) or animals (predation) – by describing how energy is consumed and transferred across various energy channels within food webs (Barnes *et al*. 2018). For instance, the energy channel through herbivores is associated with plant biomass consumption and production (Barnes *et al*. 2020); energy flux through decomposers is linked to nutrient cycling and storage (Potapov 2022); energy flux through bacteria and fungi have been proposed to represent relatively fast and slow pathways of energy and nutrient turnover, respectively (Moore *et al*. 2005); and that through predators can be related to the top-down control of the whole food web or a specific energy channel (Barnes *et al*. 2018; Potapov 2022). Odum’s predicted shift from grazing- to detritus-dominated food chains can thus be examined as a shift from herbivory-based (‘green’) to detritus-based (‘brown’) energy channels (Odum 1969; Wolkovich *et al*. 2014). If energy channels are mainly controlled by resource supply, changes in each channel should track changes in its basal resource pool along succession (Fig. 1).

**Fig. 1.**
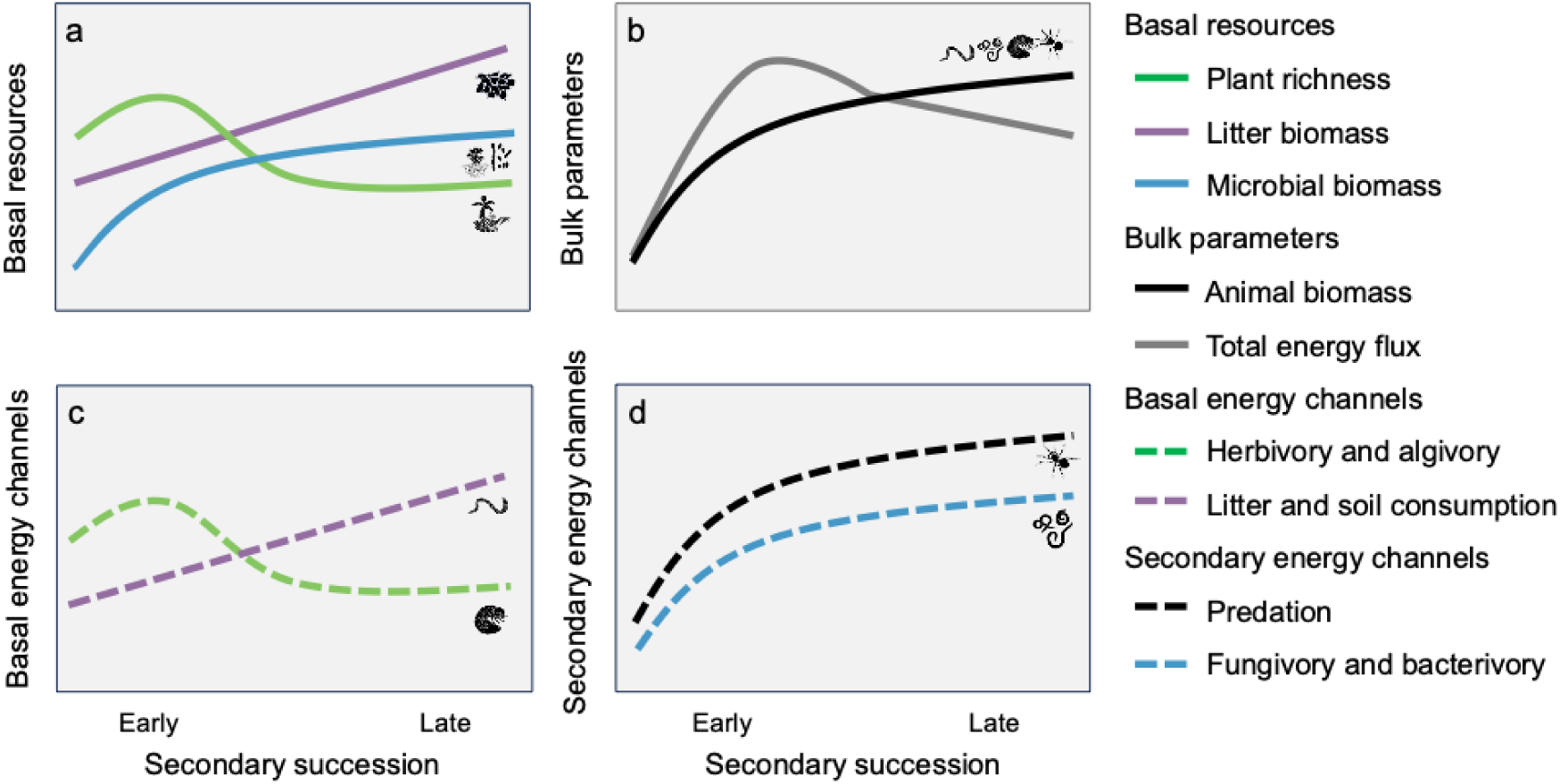
Predicted changes in soil food web energetics along secondary succession, based on Odum’s theory of ecosystem development. (Odum 1969). Basal resources (**a**), soil food-web bulk parameters (**b**), soil food-web basal energy channels (**c**), and soil food-web secondary energy channels (**d**). Trends in plant richness and litter biomass (**a**) are based on published data from the study region (Zhang *et al*. 2023); microbial (basal resource) and animal biomass are predicted to increase (Frouz *et al*. 2013; Odum 1969), whereas total energy flux peaks early and then declines, resulting in an increasing biomass-to-energy flux ratio (B/E) (**b**). Energy is predicted to shift from herbivory- to detritus-based channels (**c**), with each channel tracking its corresponding resource proxy (H1): herbivory follows plant richness, litter and soil consumption follow litter biomass, microbivory follows microbial biomass, and predation follows animal biomass (**d**). Early-successional stages: grassland and shrubland; late-successional stages: secondary and primary forests. Animal silhouettes were drawn by Svenja Meyer.

To understand soil food web functioning, we need to consider the complex size structure of these systems. Micro-food webs composed of microfauna (e.g., nematodes) are characterized by rapid turnover and reliance on pulsed energy inputs, while macro-food webs comprised of meso- and macrofauna (e.g., microarthropods, macroarthropods and earthworms) are more stable and less localised due to slower reproduction and greater mobility of the organisms (Brose *et al*. 2017; Potapov *et al*. 2021). Soil micro-food webs are therefore expected to respond more strongly to environmental changes and to be more variable in time and space (Gallego-Zamorano *et al*. 2023), whereas macro-food webs, whose members feed on a wide range of basal resources, should respond more slowly; they also contribute substantially to terrestrial arthropod diversity and ecosystem functions (Potapov *et al*. 2019b, 2021). The coupling of compartments with fast and slow dynamics has been proposed to stabilise soil food webs (Potapov *et al*. 2021; Rooney *et al*. 2006). However, whether micro- and macro-food webs change in parallel or diverge during vegetation succession remains unclear.

Here, we examined changes in the energetic structure of soil micro- and macro-food webs along secondary succession (Fang *et al*. 2024; Zhang *et al*. 2023) and explored the relationships between abiotic soil variables, basal resources, animal richness, and energy fluxes. Because succession is shaped not only by natural processes but also by human activities (Poorter *et al*. 2024), we studied a natural successional sequence (grassland, shrubland, secondary birch forest and primary forest) and compared the secondary birch forest with spruce and larch plantations of similar age, which are widespread in the region (Liu *et al*. 2002). Following Odum (Odum 1969), we hypothesised that, as organic matter and biomass accumulate along natural succession (Zhang *et al*. 2023), B/E of consumer communities would increase and energy would shift from herbivory- to detritus-based channels (Fig. 1). Specifically, we expected that trends in energy channels would track their corresponding resource proxies (i.e., herbivory, detritivory, microbivory, and predation would follow changes in plant richness, litter biomass, microbial biomass, and animal biomass, respectively) (H1). In particular, we expected herbivory to increase with rising plant richness early in succession and to decline as plant richness decreases later; detritivory to increase progressively with accumulating litter biomass; and microbivory and predation to rise with microbial and animal biomass, respectively, before plateauing towards primary forest. In addition, we hypothesised that, along vegetation succession, total energy flux in soil micro-food webs would change more than in macro-food webs (H2), reflecting the faster turnover and higher environmental sensitivity of small-bodied taxa (e.g., nematodes) than of larger soil animals (Potapov *et al*. 2021).

**Fig. 2.**
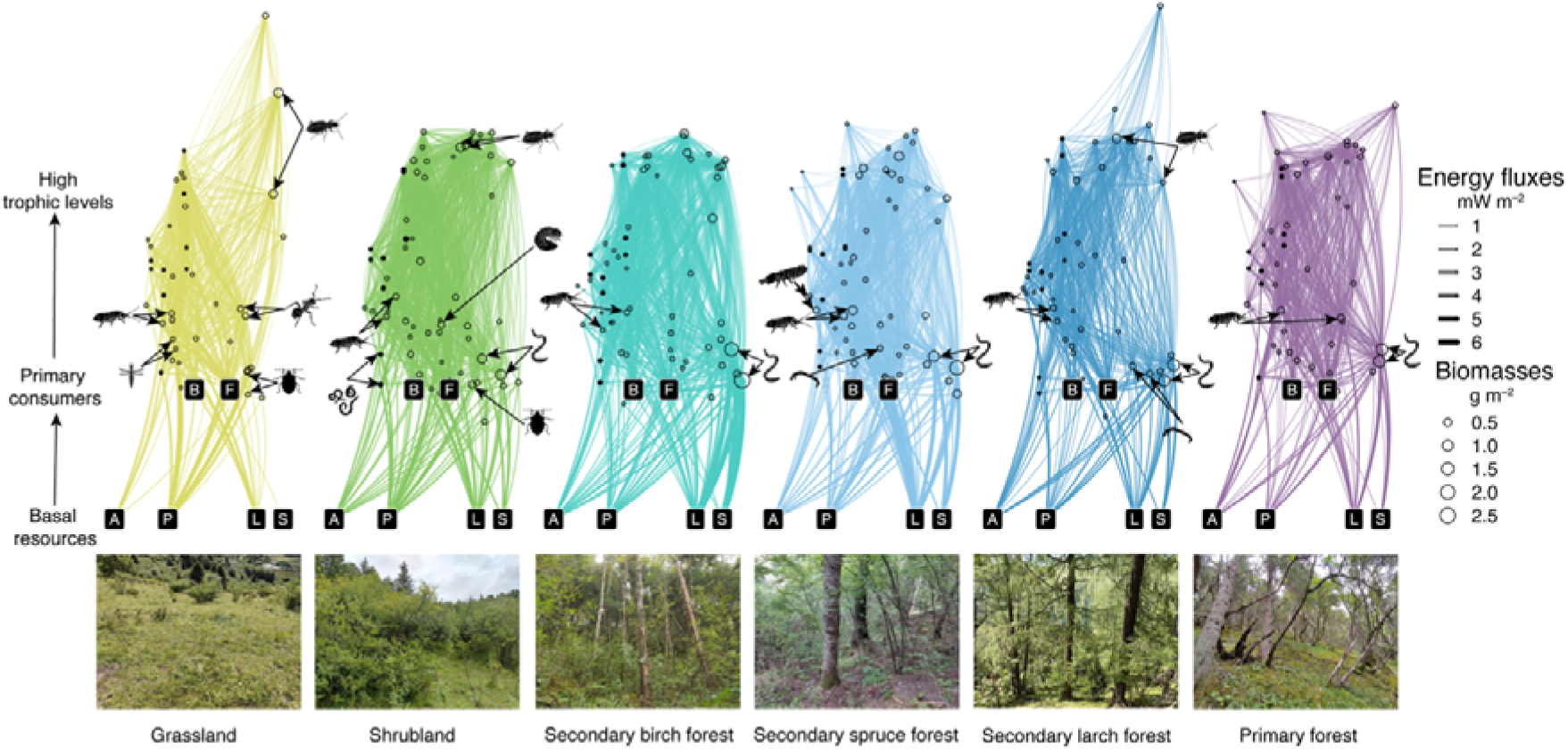
Structure of reconstructed soil food webs along secondary succession (n = 6 plots per ecosystem). Each food web shows mean values across the six plots of an ecosystem; photographs below show the corresponding ecosystems. Consumer nodes are represented by trophic guilds and are arranged by trophic level (y-axis). Resources are shown as black square labels along the x-axis (A: algae; P: living plants; L: plant litter; S: soil organic matter; F: fungi; B: bacteria). The sizes of consumer nodes are proportional to their respective biomasses, with solid black circles representing soil micro-food webs and dotted black circles representing soil macro-food webs. Average energy fluxes, quantified in mW m^−2^ (represented by line thickness), are represented by the connecting lines in the soil food web diagram. Colours denote successional stages (fluorescent green, grassland; green, shrubland; teal, secondary birch forest; light blue, secondary spruce forest; dark blue, secondary larch forest; purple, primary forest). Animal silhouettes show taxa contributing over 5% to the total energy flux in each ecosystem: Collembola (detritivores), Formicidae, Thysanoptera, Coleoptera (predators), and Hemiptera (Heteroptera herbivores) in grassland; Collembola (detritivores), Lumbricina, Coleoptera (predators and herbivores), Hemiptera (Auchenorrhyncha), and Nematoda (plant feeders) in shrubland; Lumbricina and Collembola (detritivores) in secondary birch forest; Collembola (detritivores and Onychiuroidea), Lumbricina, and Diptera (phytosaprophagous) in the spruce plantation; Collembola (detritivores), Diptera (phytosaprophagous), Coleoptera (predators) and Lumbricina in the larch plantation; Collembola (detritivores) and Lumbricina in primary forest. Animal silhouettes were drawn by Svenja Meyer. Photo credit: Ajuan Zhang.

## RESULTS

### Increase in biomass-to-energy flux ratio

Animal biomass increased along the secondary succession (linear contrast, *P* = 0.019) and peaked in the secondary birch forest (11.4 ± 3.6 [mean ± SE] g fresh biomass m^−2^, quadratic contrast, *P* = 0.030), 10.4 and 4.0 times higher than in early-successional grassland (1.1 ± 0.1 g m^−2^) and shrubland (2.9 ± 0.7 g m^−2^), respectively (Fig. 3 and Table S6). Total energy flux declined along the succession (linear contrast, *P* = 0.017), although this decline depended on the temperature differences among ecosystems (Fig. S12); it was highest in the spruce plantation (57.2 ± 7.2 mW m^−2^) and lowest in the larch plantation (21.0 ± 5.0 mW m^−2^; vs. birch forest, *P* = 0.019) and primary forest (20.1 ± 2.8 mW m^−2^) (Fig. 3 and Table S6). Consequently, B/E ratio increased (linear contrast, *P* = 0.004), peaking in the secondary birch forest (0.24 ± 0.04), 10.4 and 4.3 times higher than in grassland (0.023 ± 0.002) and shrubland (0.056 ± 0.007), respectively. Animal richness showed no significant successional change (all *P* ≥ 0.063). Changes in energy flux within size compartments generally mirrored the decline in total energy flux along secondary succession (linear contrast, *P* = 0.041 and 0.027 for the macro and micro compartments, respectively; both dependent on temperature differences, Fig. S12), with a more pronounced change in the micro compartment (85.7% reduction, from the maximum of 13.9 ± 4.0 mW m^−2^ in shrubland to the minimum of 2.0 ± 0.2 mW m^−2^ in primary forest) than in the macro compartment (58.1% reduction, from 43.3 ± 4.4 mW m⁻^2^ in grassland to 18.1 ± 2.9 mW m⁻^2^ in primary forest; ecosystem × food web interaction, *P* = 0.056; Fig. 4a and Table S7).

**Fig. 3.**
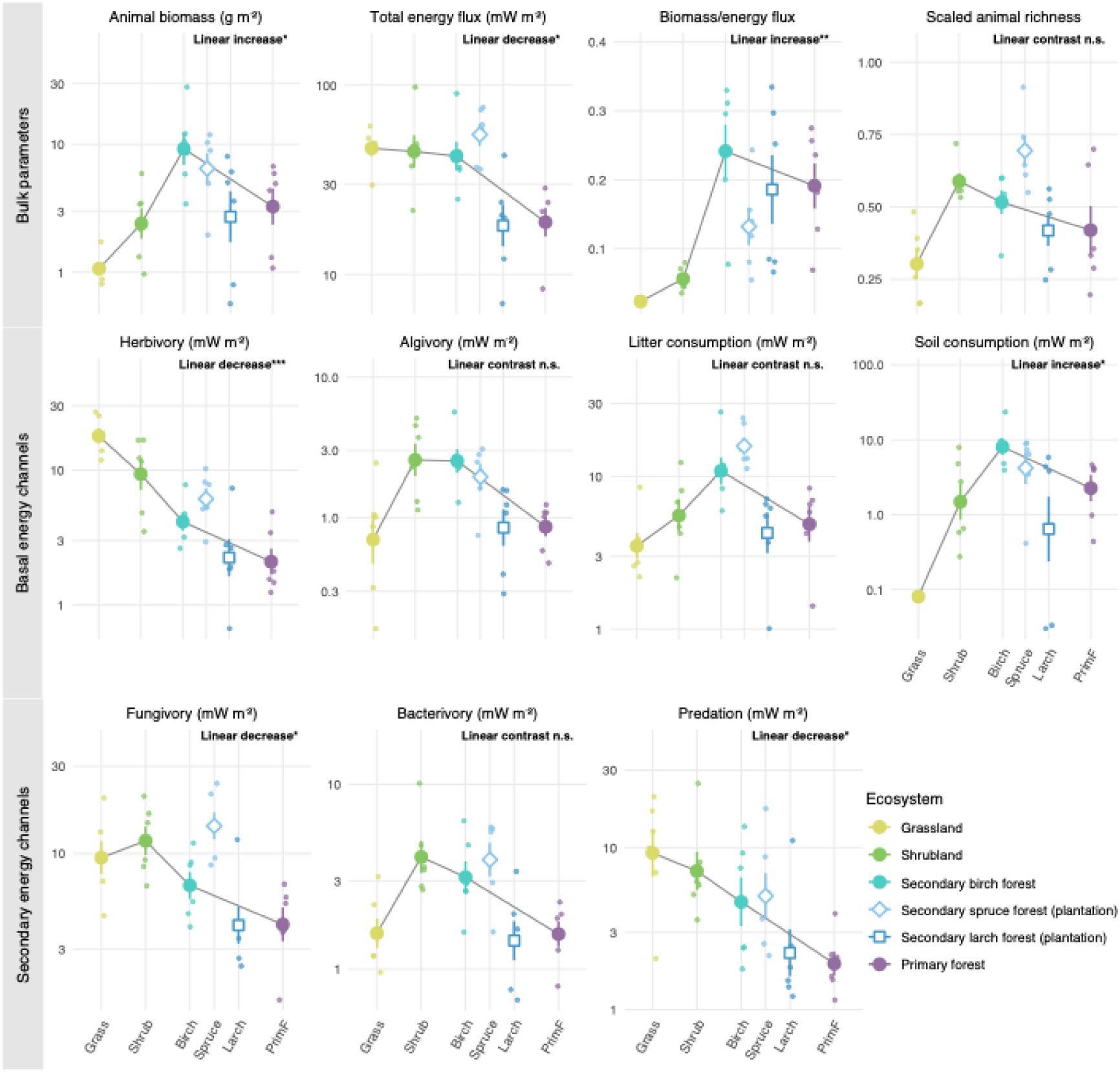
Changes in bulk parameters and energy channels of bulk soil food webs along secondary vegetation succession (n = 6 plots per ecosystem). Small points represent individual plots, and large symbols with error bars represent means ± SE. Grass: grassland; Shrub: shrubland; Birch: secondary birch forest; Spruce and Larch: spruce and larch plantations; PrimF: primary forest. Filled circles connected by lines show the natural successional sequence (grassland – shrubland – secondary birch forest – primary forest); open symbols show the spruce (diamonds) and larch (squares) plantations established in the same period as the birch forest, which are not connected to the successional sequence. Colours denote ecosystems (yellow-green, grassland; green, shrubland; teal, secondary birch forest; light blue, spruce plantation; dark blue, larch plantation; purple, primary forest). Units for each parameter are given in brackets. Labels in each panel give the linear contrast along the natural successional sequence (\**P* < 0.05, \*\**P* < 0.01, \*\*\**P* < 0.001; n.s., not significant), based on linear mixed-effects models with ecosystem and river basin as fixed effects and site as a random effect. All contrasts, including comparisons of plantations with secondary birch forest, are given in Table S6.

**Fig. 4.**
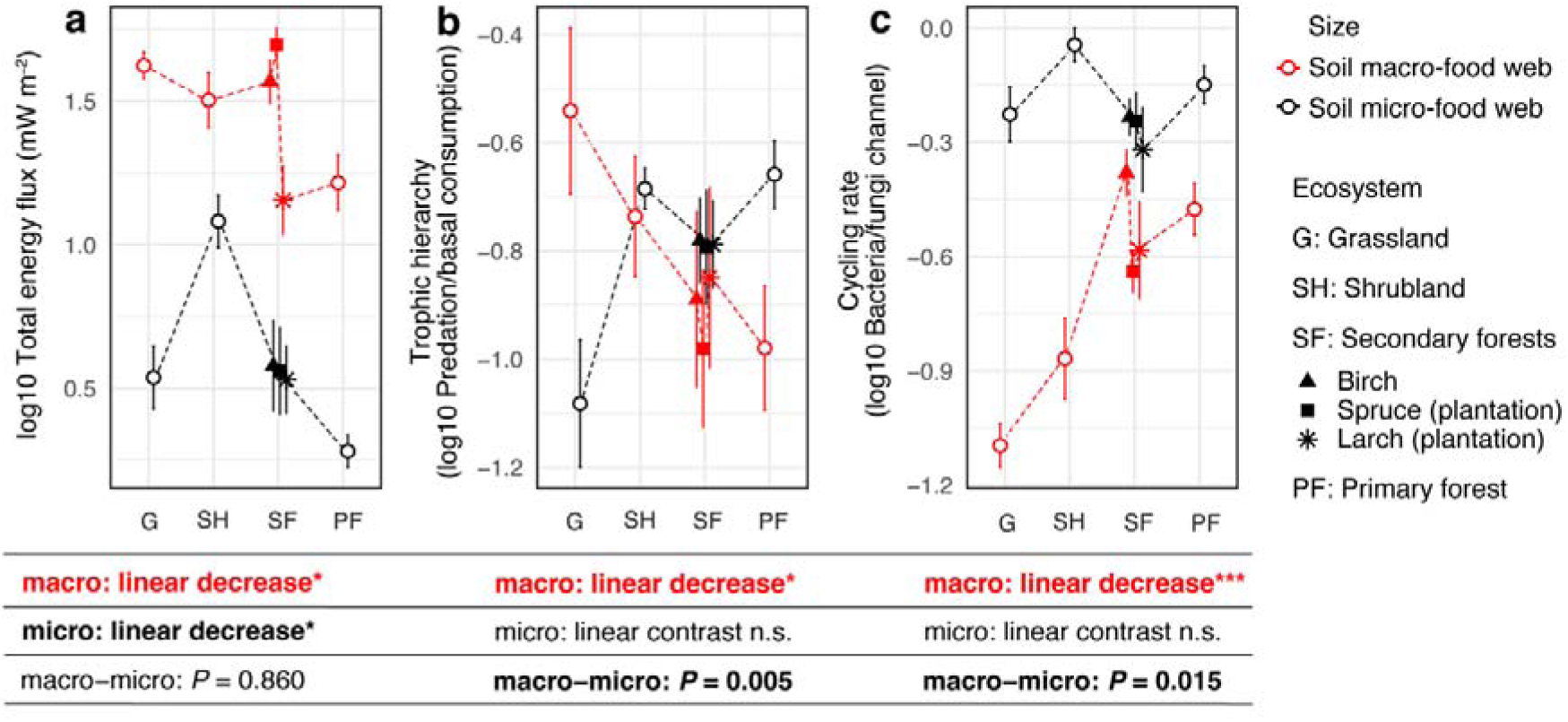
Changes in soil micro- and macro-food web indicators along secondary vegetation succession (n = 6 plots per ecosystem). **a–c**, Total energy flux (**a**); trophic hierarchy (**b**); cycling rate (**c**). The trophic hierarchy is calculated as the ratio of all the energy fluxes from ‘predation’ to all the energy fluxes from ‘consumption of basal resources’. The cycling rate is calculated as the ratio of all outgoing fluxes from bacteria to all outgoing fluxes from fungi. All variables were log10-transformed. Points represent means and error bars indicate standard errors. Red colours represent soil macro-food webs and black colours represent soil micro-food webs. G: grassland; SH: shrubland; SF: secondary forests including birch (triangles), spruce (squares, plantation) and larch (asterisks, plantation) forests; PF: primary forest. Text below each panel gives the linear contrast along the natural successional sequence (grassland – shrubland – secondary birch forest – primary forest) for each food web and the difference in this linear trend between macro- and micro-food webs (* *P* < 0.05, *** *P* < 0.001; n.s., not significant), based on linear mixed-effects models with ecosystem, food web, their interaction and river basin as fixed effects, and site, site × food web and plot as random effects. All contrasts are given in Table S7.

### Change of energy distribution

Energy channels (i.e., energy fluxes originating from specific resources) responded differently to succession (Fig. 3 and Table S6). Herbivory declined along succession (linear contrast, *P* < 0.001) and was 2.8 to 7.8 times lower in secondary birch forest, both plantations and primary forests, than in grassland (18.8 ± 2.5 mW m^−2^). Fungivory also declined (linear contrast, *P* = 0.049), from 12.7 ± 2.2 mW m^−2^ in shrubland to 4.8 ± 0.8 mW m^−2^ in primary forest, whereas soil consumption increased (linear contrast, *P* = 0.021) from < 0.1 mW m^−2^ in grassland to a peak in the secondary birch forest (9.6 ± 2.9 mW m^−2^; quadratic contrast, *P* = 0.033). Algivory, litter consumption and bacterivory showed hump-shaped patterns along the natural successional sequence (quadratic contrast, *P* = 0.007, 0.039 and 0.011, respectively), peaking in shrubland (algivory, 3.0 ± 0.7 mW m^−2^; bacterivory, 4.6 ± 1.2 mW m^−2^) or the secondary birch forest (litter consumption, 12.2 ± 3.0 mW m^−2^) and being lowest in grassland and primary forest. Compared with the secondary birch forest, the larch plantation had lower algivory, litter consumption, soil consumption and bacterivory (*P* = 0.034–0.045), whereas the spruce plantation did not differ in any energy channel (all *P* ≥ 0.10).

### Size class-specific changes in predation

Predation was highest in grassland (11.5 ± 2.7 mW m^−2^) and declined along succession (linear contrast, *P* = 0.029), being 72% to 82% lower in the larch plantation and primary forest (3.2 ± 1.6 and 2.1 ± 0.4 mW m^−2^, respectively; Fig. 3 and Table S6). The predation-to-basal consumption ratio, as an energetic proxy for top-down control, decreased along succession in macro-food webs (linear contrast, *P* = 0.049) but tended to increase in micro-food webs (*P* = 0.100); these trends differed significantly between food webs (difference in linear contrast, *P* = 0.005; ecosystem × food web interaction, *P* = 0.040; Fig. 4b and Table S7), whereas the overall ratio did not change significantly throughout succession (all *P* ≥ 0.12; Fig. S11 and Table S6).

### Shift in carbon cycling

The herbivory-to-detritivory ratio (representing the balance between ‘green’, i.e., herbivory-dominated, and ‘brown’, i.e., detritivory-dominated, energy pathways) (Wolkovich *et al*. 2014) was highest in grassland (5.9 ± 1.1) and declined along succession (linear contrast, *P* < 0.001), being 89% to 94% lower in secondary birch forest, both plantations and primary forest (Fig. S11 and Table S6). The bacteria-to-fungi energy flux ratio (a proposed proxy for ‘fast’ versus ‘slow’ energy turnover) increased along succession (linear contrast, *P* = 0.040), peaking in the secondary birch forest (0.48 ± 0.04; Fig. S11 and Table S6). Here, again, trends differed between size compartments (difference in linear contrast, *P* = 0.015; ecosystem × food web interaction, *P* = 0.063; Fig. 4c and Table S7). In soil macro-food webs, the bacterivory-to-fungivory ratio increased along succession (linear contrast, *P* < 0.001), from grassland (0.08 ± 0.01) and shrubland (0.16 ± 0.04) to the secondary birch forest (0.44 ± 0.06) and primary forest (0.35 ± 0.05). In contrast, the ratio in soil micro-food webs showed no successional change (all *P* ≥ 0.58). Additionally, the faeces production-to-soil organic matter consumption ratio (a potential proxy for animal contribution to soil organic matter sequestration) was highest in grassland (27.3 ± 12.4; high production) and 71% to 98% lower in the other ecosystems, with the lowest value in the secondary birch forest (0.6 ± 0.3; high consumption), but these differences were not significant (all *P* ≥ 0.25; Fig. S11 and Table S6).

### Environmental drivers

We used partial least squares path models (PLS-PM) to explore the effects of multitrophic animal richness, basal resources (plant richness, litter quantity and quality, and microbial biomass), and abiotic soil variables on energy flux across successional stages (Fig. 5). Total energy flux was positively related to animal taxonomic richness (*β* = 0.56) and microbial biomass (*β* = 0.37) and negatively to litter quantity (*β* = −0.40); the overall effect of succession on energy flux was negative (*β* = −0.47). Plant richness, litter quantity and abiotic soil variables also affected total energy flux indirectly via animal richness (*β* = −0.39, 0.43 and 0.52, respectively), and litter quality via microbial biomass (*β* = −0.53). Energy flux in soil macro-food webs showed the same pattern as total energy flux, whereas energy flux in micro-food webs was positively associated only with litter quality. Among energy channels, animal richness was positively associated with all channels, and herbivory and predation were additionally positively associated with litter quality and microbial biomass.

**Fig. 5.**
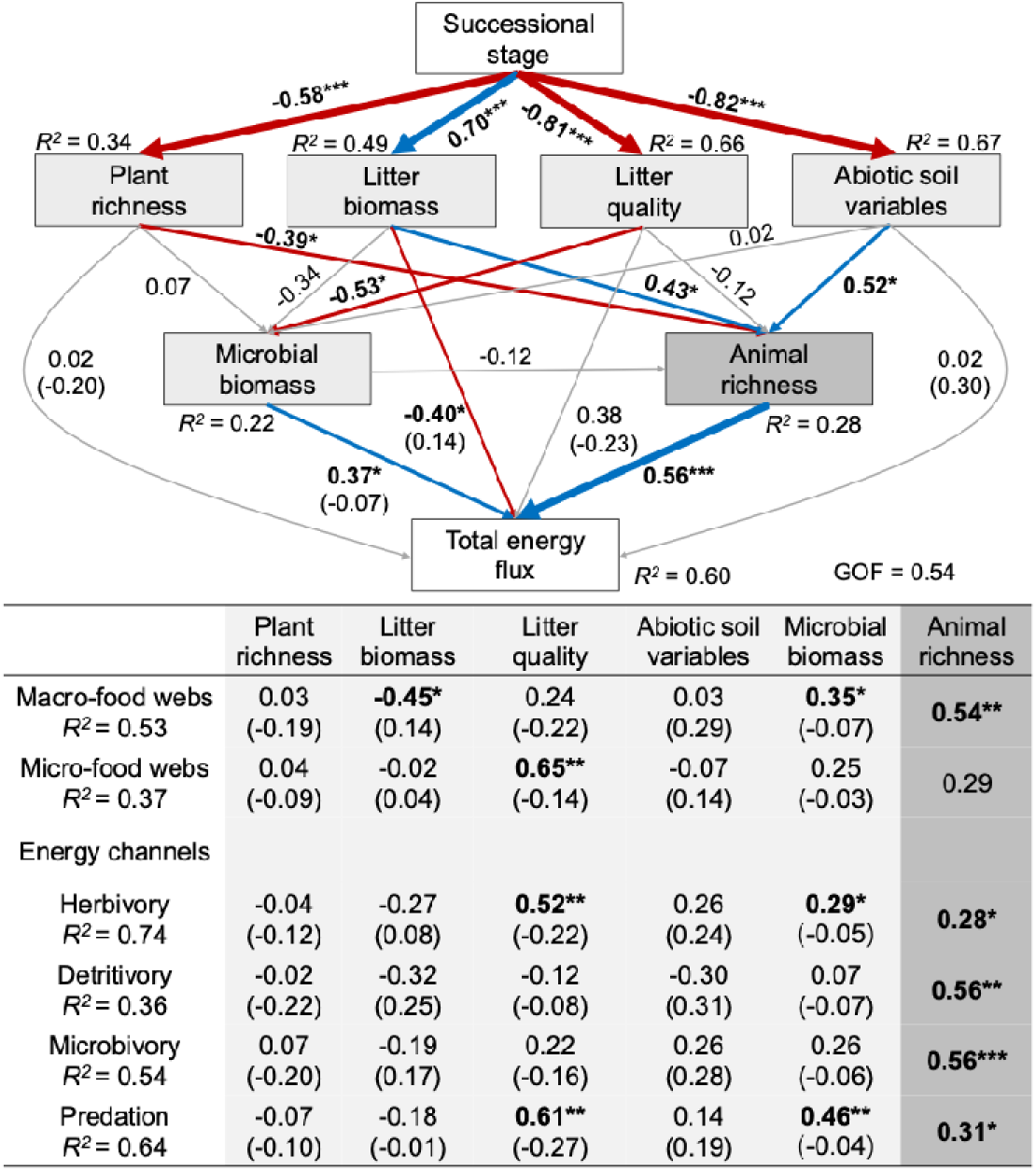
Partial least squares path models (PLS-PM) illustrating the relationships between secondary vegetation succession, basal resources, multitrophic animal richness and the total energy flux, energy flux of soil macro- and micro-food webs, and individual energy channels (n = 36 plots). Succession stage was coded as a numerical variable (grassland = 1, shrubland = 2, secondary birch forest and spruce and larch plantations = 3, primary forest = 4); results restricted to the natural successional sequence are given in Table S8. Numbers adjoining the arrows indicate standardized path coefficients, and the width of the arrows is proportional to the strength of the association. In the lower panel, values are standardized direct path coefficients of each predictor on the respective energy flux, with indirect effects in parentheses. Paths were considered significant when their 95% bootstrap confidence intervals (1000 resamples) excluded zero; asterisks give P values (* *P* < 0.05, ** *P* < 0.01, *** *P* < 0.001). Blue arrows represent significant positive relationships, red arrows represent significant negative relationships, and grey arrows represent non-significant relationships. *R*² values indicate the proportion of variance of each endogenous variable explained by the model. Litter quality was represented by litter C/N ratio (reversed) and cellulose concentration (reversed), and abiotic soil variables by soil pH, soil temperature and soil C/N ratio (reversed). Energy fluxes were log10-transformed. GOF, goodness of fit. Light grey represents basal resources and dark grey represents soil fauna.

### Sensitivity analyses

Most results were robust to the temperatures used to correct metabolic rates (Fig. S12). The increase in B/E ratio, the declines in herbivory and in the herbivory-to-detritivory ratio, the increases in soil consumption and in the bacterivory-to-fungivory ratio, the hump-shaped patterns of algivory, litter consumption, bacterivory and micro-food-web energy flux, and the differences in predation-to-basal consumption and bacterivory-to-fungivory ratios between macro- and micro-food webs remained significant when temperature differences among ecosystems were halved or removed. In contrast, the declines in total energy flux (including those of macro- and micro-food webs), fungivory and predation along succession were no longer significant, indicating that they mainly reflected the higher soil temperatures in grassland and shrubland. Varying the weight of auxiliary food sources changed absolute energy fluxes, particularly litter and soil consumption, but the relative ordering of ecosystems remained largely consistent (Fig. S13). Decomposition by taxon showed that earthworms, which were absent from grassland and had the lowest energy flux per unit biomass of all taxa, made up 45–86% of animal biomass in shrubland and forests. Without earthworms, B/E still increased under baseline temperatures (linear contrast, *P* < 0.001) but not under a common temperature (*P* = 0.988; Fig. S14).

## DISCUSSION

Our study reveals shifts in energy turnover and distribution, describing the functional development of soil food webs along secondary succession. In line with Odum’s framework of ecosystem development (Odum 1969), the ratio of animal biomass to energy flux in soil food webs increased linearly from grassland to primary forest, indicating longer energy turnover times in more mature ecosystems. This pattern matches the increase in biomass supported per unit energy flow that Odum predicted for mature ecosystems, and reflects the accumulation of consumer biomass in soil (Odum 1969). We further show a transition from a herbivory-dominated ‘green functional state’ to a detritivory-dominated ‘brown functional state’ of soil food webs, underscoring a fundamental shift in energy channelling in the ecosystem. Resolving energy fluxes separately for the macro- and micro-food webs further allowed us to show changes in the energy fluxes at higher trophic levels. In particular, the bacterivory-to-fungivory ratio increased along succession in soil macro-food webs but not in micro-food webs, and the ratio of predation to basal consumption followed opposite trends in the two compartments, increasing in micro- relative to macro-food webs. Moreover, tree plantations established at the same successional stage did not consistently match secondary birch forest: the spruce plantation was similar to it, whereas the larch plantation had lower animal biomass and energy fluxes. Overall, our results extend Odum’s framework to soil food webs by showing systematic shifts in energy channelling across trophic levels in soil food webs (Odum 1969), and provide a methodological approach for studying the functional development of soil food webs.

### Energy turnover and energy flux along succession

Along secondary succession, energy flux per unit animal biomass declined (that is, an increase in B/E) (Fig. 3), consistent with slower energy turnover in more mature ecosystems (Buzhdygan *et al*. 2020; Odum 1969). This increase was robust to the assumed temperature differences and diet compositions (Section 3.6), and arose because animal biomass increased while total energy flux did not. Decomposition by taxon showed that it was mainly driven by earthworms, which were absent from grassland but made up 45% of animal biomass in shrubland and 67–86% in forests (Figs. S4 and S14). Owing to their large body size and low metabolic rates, earthworms had the lowest energy flux per unit biomass of all taxa; after excluding earthworms and removing temperature differences, B/E no longer increased (Fig. S14). As earthworms were hand-sorted, their absence from grassland is unlikely to be a sampling artefact and more likely reflects the warm, dry soils and low litter input of the open grassland (Figs. S2 and S3), because earthworms are sensitive to desiccation and heat and feed on litter and soil organic matter (Potapov *et al*. 2022). Conversely, small, fast-turnover insects such as ants, hemipterans and thrips, which made up about 19% of animal biomass in grassland, nearly disappeared in forests. This replacement of small insects by large detritivores parallels the shift from green to brown energy channelling. These results support Odum’s prediction that biomass accumulates while energy turnover declines as ecosystems develop (Odum 1969), and show that in soil animal communities this pattern is largely mediated by the establishment of earthworms.

Total energy flux declined in both macro- and micro-food webs (Figs. 3 and 4a), consistent with the cooler soils under closed canopies, as metabolic rates increase with temperature (Brown *et al*. 2004). In soil macro-food webs, this decline was accompanied by a decrease in predation relative to basal consumption (Fig. 4b), although this trend was sensitive to the temperature correction (Fig. S12). In soil micro-food webs, by contrast, this ratio tended to increase, and its trend differed significantly from that in macro-food webs (Fig. 4b; Table S7), suggesting that a growing share of energy flowed to predatory nematodes in more mature ecosystems. This pattern aligns with studies on nematode community succession, where mean body size and trophic level were reported to increase as ecosystems mature (Ferris *et al*. 2001; Zhang *et al*. 2025b). Moreover, energy flux in micro-food webs was hump-shaped, peaking in shrubland, which paralleled the pattern of bacterivory (Figs. 3 and 4a). Together with the synchronous shift of plants and nematodes from small, fast-growing to larger, slower-growing organisms during natural restoration (Zhang *et al*. 2025b), this suggests that nematodes respond sensitively to vegetation succession.

### Transition from green to brown functional state

Our findings reveal a strong decrease in the herbivory-to-detritivory flux ratio along the succession (Fig. S11), indicating a transition of soil food webs from a herbivory-dominated ‘green functional state’ in grassland to a detritivory-dominated ‘brown functional state’ in forests. This transition resulted from two parallel changes. First, herbivory declined by an order of magnitude (from 18.8 to 2.4 mW m^−2^ between grassland and primary forest; Fig. 3), as herbivorous insects such as hemipterans, thrips and ants, which accounted for 82% of herbivory in grassland, nearly disappeared under forest canopies. Second, soil consumption increased (Fig. 3), reflecting the increase in earthworm biomass, which accounted for 90−98% of this flux in shrubland and forests. Litter consumption, in contrast, did not increase linearly but peaked in the secondary birch forest. As succession proceeded, litter biomass and the C/N ratios of litter and soil increased (Figs. S2 and S3), indicating accumulation of detrital resources of lower quality (Hooper *et al*. 2000). Rather than increasing litter consumption, this accumulation appears to have favoured soil-feeding earthworms, shifting the basal resource of soil food webs towards soil organic matter. In addition, maturation is accompanied by changes in belowground plant traits, such as denser and deeper root systems and mycorrhizal associations that retain mineral nutrients in the rhizosphere (Poorter *et al*. 2023); such tighter internal nutrient cycling may further favour decomposer-driven fluxes (Morriën *et al*. 2017; Zou *et al*. 2016). In summary, the transition from green to brown functional states highlights how principal energy pathways change during succession. Odum described the shift from grazing- to detritus-based food chains at the whole-ecosystem level, in which soil food webs are usually viewed as the detrital compartment (Odum 1969). We show that this shift also occurs within soil food webs, driven by the replacement of herbivorous insects by soil-feeding earthworms, which builds the brown state on soil organic matter rather than on litter. Because the same replacement also increased B/E, two attributes that Odum listed separately, the type of food chain and the biomass supported per unit energy flow, are linked through a single change in soil community composition. These findings provide deeper insights into the specific role of soil consumers in regulating of nutrient cycling and driving the transition to a ‘brown’ state in mature ecosystems.

### Predation and size compartmentalisation

We initially hypothesised that predation would increase along with animal biomass during succession; however, despite the increase in animal biomass, predation declined (Fig. 3). Because the overall ratio of predation to basal consumption did not change significantly (Fig. S11), this decline largely paralleled the decrease in total energy flux. However, the two size compartments showed diverging trends: the ratio decreased in macro-food webs but tended to increase in micro-food webs (Fig. 4b). The decrease in macro-food webs was driven by the rise of earthworms: after excluding them, the ratio no longer declined (Fig. S14). Large detritivores such as earthworms typically have fewer connections to predators (Potapov *et al*. 2019b, 2021) and can act as energy sinks, retaining energy within their large bodies rather than passing it to higher trophic levels (Potapov *et al*. 2019b; Schwarzmüller *et al*. 2015). Macro-food webs thus became more bottom-heavy as consumer biomass increased, consistent with the sublinear scaling of predator to prey biomass reported across ecosystems (Hatton *et al*. 2015). Conversely, the increase in this ratio in micro-food webs may reflect a growing share of large nematode predators as ecosystems mature, consistent with nematode succession studies (Ferris *et al*. 2001). This contrast reflects the size-structured nature of soil micro-food webs, where larger organisms typically function as predators, whereas in macro-food webs larger organisms often function as basal consumers (Potapov *et al*. 2021). These patterns show that organism size and compartmentalisation must be considered when studying energy flux and ecosystem development and support size spectrum theory as a framework for understanding how energy constraints shape community structure (Brose *et al*. 2017; Potapov *et al*. 2019a, 2021).

### Bacterial and fungal energy channels

The bacterivory-to-fungivory ratio increased along succession (Fig. S11), mainly because fungivory declined while bacterivory showed no linear trend (Fig. 3). Assuming that bacterial and fungal energy channels are partly independent (Pollierer *et al*. 2012), bacteria and their consumers form a fast cycle, whereas fungi and their consumers turn over more slowly (Hunt *et al*. 1987; Rooney *et al*. 2006; de Ruiter *et al*. 1995). The bacterial energy channel, which primarily processes labile organic matter, typically dominates in fertile, productive ecosystems such as grasslands, whereas the fungal channel prevails in less fertile ecosystems such as forests, where litter is rich in lignin and structural compounds but poor in nitrogen (Wardle *et al*. 2004a). Accordingly, soil food webs are expected to shift from bacterial to fungal dominance during succession (Frouz *et al*. 2013; Wardle *et al*. 1995). Our results contradict this expectation, even though litter and soil C/N ratios increased (Figs. S2 and S3). The ratio increased only in macro-food webs and remained unchanged in micro-food webs (Fig. 4c), indicating that soil macrofauna drove the overall trend. In our reconstruction, this reflected the replacement of fungus-feeding insects, notably thrips, which accounted for 39% of fungivory in grassland, by earthworms, which assimilate bacterial carbon from older soil organic matter (Hyodo *et al*. 2008; Potapov *et al*. 2024). Through bioturbation and litter-soil mixing, earthworms can shift energy channels towards bacterial dominance (Frouz *et al*. 2008, 2013; Liu *et al*. 2019), potentially accelerating the depletion of older carbon stocks (De Vries *et al*. 2013). However, earthworms play a dual role in carbon cycling: they incorporate organic material into the soil through faeces production, enhancing microbial activity, but also deplete soil organic matter by consuming it (Joly *et al*. 2020; Potapov *et al*. 2024). The establishment of earthworms during succession may therefore accelerate soil carbon turnover through the bacterial channel, but its net effect on soil carbon storage remains to be tested.

### Biotic and abiotic drivers of energy flux

The effects of secondary succession on energy flux were primarily mediated by animal richness and basal resources (microbial biomass, litter quantity and quality), and abiotic variables (Fig. 5). Animal richness was the strongest predictor of total energy flux and was positively associated with all four energy channels, which aligns with the widely recognised relationship between multitrophic biodiversity and energy flux often found in ecological communities (Barnes *et al*. 2014, 2018). This connection may stem from niche or trophic complementarity, where greater consumer diversity reduces resource competition, thereby enhancing total energy transfer to consumers within food webs (Barnes *et al*. 2018; Poisot *et al*. 2013). Furthermore, the observed positive correlation between microbial biomass and both total energy flux and energy flux from predation and herbivory channels suggests that increased microbial biomass may further boost energy availability for consumers, reinforcing the overall energy flow within the ecosystem (Schramski *et al*. 2015) and supporting earlier published evidence that soil food webs are bottom-up controlled (Scherber *et al*. 2010). By contrast, litter quantity exhibited a negative association with the total energy flux and with energy fluxes in macro-food webs. This effect is rather enigmatic and may be attributed to the accumulation of stable, unpalatable plant residues with low decomposability along succession (Wardle *et al*. 2004b), which subsequently affects the resource quality available to soil fauna, thereby contributing to a negative feedback on energy flux. In summary, contrary to our expectations, the variations in energy channels did not always align consistently with their corresponding resource changes (i.e., herbivory, detritivory, microbivory, and predation did not uniformly follow shifts in plant richness, litter biomass, microbial biomass, and animal biomass, respectively). This discrepancy implies that factors beyond simple resource availability may regulate energy flux in these channels, suggesting a more complex interplay of biotic and abiotic factors (and not simply the bottom-up control) influencing ecosystem functioning throughout succession. Therefore, our findings underscore the value of adopting the energy flux approach, which integrates metabolic rates, assimilation efficiency, and trophic interactions, to provide a more comprehensive understanding of multitrophic biodiversity–ecosystem functioning relationships (Pollierer *et al*. 2023).

### Limitations and perspectives

Our approach has limitations that also open avenues for future research. Like most studies of succession, our design substitutes space for time (Walker *et al*. 2010); we reduced confounding by replicating each ecosystem in two river basins, treating sites as experimental units and analysing plantations separately from the natural sequence. Energy fluxes were reconstructed rather than measured directly, but the central patterns, the increase in B/E and the transition from green to brown energy channelling, were robust to assumptions on temperature and diet composition. Because these patterns hinge on earthworms, stable isotope or fatty acid analysis of earthworms’ diets would further constrain the reconstruction. Finally, including protists (Geisen 2016) and repeated sampling across seasons would extend our view of micro-food web energy flux and its temporal dynamics.

## CONCLUSIONS

Our study quantitatively showed that soil food webs shift from fast to slow energy turnover and from green to brown energy channelling along secondary succession, thereby emphasizing the pivotal role of soil community composition in ecosystem energetics. Both shifts were largely driven by a single change in community composition: the replacement of small, fast-turnover herbivorous insects by large, soil-feeding earthworms. Our findings revealed that size compartments of soil food webs diverged along the successional gradient: macro-sized compartments shifted towards bottom-heavy, bacterial-based food webs, while micro-sized compartments evolve towards top-heavy, fungal- and bacterial-based food webs. This dichotomy underscores a dual mechanism of soil functioning where one compartment (i.e., soil macro-food web) efficiently sequesters biomass, while the nested food web (i.e., soil micro-food web) maintains rapid nutrient cycling. Tree plantations did not consistently match secondary birch forest of the same stage, as the larch plantation supported lower animal biomass. Together, these results show that the functional development of soil food webs can hinge on the establishment of a few key taxa rather than on changes in overall diversity. Future studies should test whether these patterns hold in other ecosystems and regions, and in primary succession. Examining both aboveground and belowground energy distribution during succession could further reveal the belowground–aboveground feedbacks that drive ecosystem development (Jochum & Eisenhauer 2022). Such knowledge could inform restoration strategies by considering whether key soil taxa establish, rather than focusing on vegetation recovery alone.

## MATERIALS AND METHODS

### Study region and experimental design

This study was conducted in the Miyaluo Forest region of Lixian County, located in western Sichuan, China (102°35’E**–**102°44’E, 31°35’N–31°53’E, 3200–3700 m a.s.l.), which is situated on the eastern Tibetan Plateau (Fig. S1), a globally recognised biodiversity hotspot. Following mid-20th century deforestation that reduced primary forest cover by >50% in only 40 years (WINKLER 1996), the landscape has transitioned through a mosaic of successional ecosystems. This recovery trajectory reflects both spontaneous natural regeneration (e.g., birch-dominated secondary forests) and human-mediated interventions, particularly monoculture plantations of *Larix spp.* and *Picea asperata* established since the 1980s (Liu *et al*. 2002). In protected areas, post-deforestation succession typically follows the sequence: grassland → shrubland → secondary forest → primary forest (Liu *et al*. 2002; Zhang *et al*. 2025a). Using a space-for-time approach (Moreno-Mateos *et al*. 2020), we selected four stages of this natural successional sequence (grassland, shrubland, secondary birch forest and primary forest). Because succession is also driven by human activities (Poorter *et al*. 2024), we additionally selected spruce and larch plantations established during the same period as the secondary birch forest (approximately 30 years old; Table S1), which were compared with the birch forest to assess whether human-driven succession differs from natural succession. The space-for-time approach assumes that the ecosystems differ mainly in successional stage; differences in forest type and management history between the plantations and the birch forest are therefore treated as a separate comparison rather than as successional change. This secondary successional gradient provides a natural experiment to quantify biodiversity-function relationships across ecosystem states (Kardol *et al*. 2006). We classified grassland and shrubland as early-successional stages and secondary and primary forests as late-successional stages (Poorter *et al*. 2023). The two river basins were approximately 6 km apart (Fig. S1) and were treated as spatial blocks, each containing one site of each of the six ecosystems (12 sites in total; 2 sites per ecosystem); the two sites of each ecosystem were 4.4–7.0 km apart. Within each site, three replicate sampling plots of 10 × 10 m were established at least 50 m apart, resulting in 36 plots. Because plots within a site share soil, microclimate and land-use history, sites rather than plots were treated as independent replicates in the statistical analyses. All samplings and measurements were conducted within these plots. Tree surveys were performed across the entire plot, while three shrub quadrats (2 × 2 m) and five herbaceous quadrats (1 × 1 m) were randomly established within each 10 × 10 m plot to survey shrub and herbaceous species, respectively. We used Flora Reipublicae Popularis Sinicae and the National Plant Specimen Resource Center (http://www.cvh.ac.cn) to identify terrestrial herbs, shrubs, and trees. Plant species richness was determined as the number of species recorded within each plot. Detailed information on the ecosystems, including forest age, location and topography, is available in Table S1.

### Litter and soil sampling and analysis

Samples were taken in August 2022. To assess the quantity and quality of litter resources, three litter subsamples (10 × 10 cm) were collected from each plot. These subsamples were combined after removing coarse woody debris and stones. Additionally, ten intact soil cores (5 cm diameter and 0–10 cm depth) were randomly collected from each plot after carefully removing litter and O horizon material; these cores were then combined into a composite sample to collect nematodes and examine abiotic soil variables. Soil bulk density was measured using three soil samples collected from each plot with a cylindrical sampler (0–10 cm soil depth; volume ca. 100 cm^3^). Samples were immediately placed on ice and transported to the laboratory for analysis. Raw soil samples were processed by carefully removing all visible stones and plant debris with tweezers, followed by sieving through a 2 mm mesh. The sieved soil samples were then thoroughly mixed and divided into three subsamples. The first subsample was stored at –20°C for microbial biomass analysis. The second subsample was used to analyse ammonium nitrogen (NH_4_^+^-N) and nitrate nitrogen (NO_3_^−^-N), while the third subsample was air-dried for soil physicochemical property analysis.

Dry litter mass (t ha^−1^) was measured by drying at 60°C for 48 h and weighing to constant weight. An elemental analyser (Vario MACRO cube, Elementar, Germany) was used to determine the total C and N contents of both litter and soil. Litter cellulose and lignin concentrations were determined using a modified acid detergent method (Van Soest & Wine 1968). Soil temperature (ST) was measured at 10 cm depth in each plot with a thermocouple during sampling; six readings were taken per plot and averaged. Plot means were used in the statistical analyses, and ecosystem means were used to temperature-correct metabolic rates in the energy flux calculations (Section 2.6). Soil pH was measured in a 1:2.5 soil:water suspension using a pH meter. Soil moisture (SM) was determined through the conventional oven drying and weighing method (105°C for 48 h). Cylindrical sampler soil samples were dried at 105°C for 48 h for gravimetric bulk density (BD). Soil total phosphorus (TP) was measured colorimetrically after digestion of soil samples in H_2_SO_4_–HClO_4_, and soil available phosphorus (AP) was measured colorimetrically after extraction of soil with HCl–NH_4_F. NH_4_^+^-N and NO_3_^−^-N were extracted with 2M KCl and subsequently measured with a continuous flow analyser (AA3, SEAL Analytical, Germany). Changes in basal resources, litter quality and abiotic soil variables along secondary succession are presented in Table S2 and Supplementary Figs. S2 and S3.

### Soil microbial biomass

Soil microbial biomass was assessed using phospholipid fatty acid (PLFA) analysis, following the protocol described by ref. (Bossio & Scow 1998). Lipids were extracted from 3 g of freeze-dried soil with a chloroform:methanol buffer (1:2:0.8, v:v). Phospholipids were then separated from nonpolar lipids and converted into fatty acid methyl esters (FAMEs) for quantification. The FAMEs were analysed using a gas chromatograph (Agilent 7890, Santa Clara, USA) equipped with a flame ionisation detector and an HP-Ultra 2 column (25.0 m × 200 μm × 0.33 μm). Identification was performed using the MIDI Sherlock Microbial Identification System (MIDI Inc.). Specific PLFAs served as biomarkers for various microbial groups: Gram-positive bacteria (iso-, anteiso-, and 10Me-branched PLFAs), Gram-negative bacteria (monounsaturated and cyclopropyl PLFAs), and fungi (16:1ω5c, 18:1ω9c, and 18:3ω6c) (Joergensen 2022; Willers *et al*. 2015). Non-specific markers included PLFAs 14:0, 16:0, 17:0, 18:0, and 20:0 (Joergensen 2022). Total microbial biomass was calculated as the sum of all identified PLFAs, while bacterial biomass was determined by summing the PLFAs associated with Gram-positive and Gram-negative bacteria.

### Soil nematodes

Nematodes were extracted from 50 g of fresh soil using modified Baermann funnels (Keith *et al*. 2006) and counted under an OLYMPUS BX53 microscope (Olympus, Tokyo, Japan). Abundance was expressed as individuals per 100 g dry soil and converted to individuals per m^2^ using soil bulk density and sampling depth (0–10 cm). We randomly identified 100 individuals (or all individuals if abundances were below 100) from each nematode sample to the genus level and assigned them to five trophic groups (bacterivores, fungivores, herbivores, omnivores, and predators) following established classifications (Yeates *et al*. 1993) and Nemaplex (http://nemaplex.ucdavis.edu); genus proportions were used to allocate total abundance among genera. Maximum body width was measured with “ToupView” software (ToupTek Photonics, 2013) for 10 individuals per genus (or fewer if abundances were below 10) and averaged (George & Lindo 2015). Fresh body mass of nematodes was estimated using published formulas based on maximum body width (Zhao *et al*. 2019), and the biomass of each trophic group was calculated as the sum of genus abundance multiplied by genus body mass (see Table S3 for formulas, list of identified taxa, feeding groups, fresh body mass and maximum body width).

### Soil macro- and meso-fauna

Soil macrofauna were collected by hand in the field from three random samples of litter (25 × 25 cm) and the underlying mineral soil layer (depth 10 cm) within each plot and preserved in 75% ethanol. Soil mesofauna were collected from three random litter (10 × 10 cm) and soil samples taken with a cylindrical corer (depth 10 cm; volume c 100 cm^3^) within each plot, placed in a black cloth bag to prevent escape, and transported to the laboratory. To account for spatial heterogeneity within plots, the three random samples of each layer were combined into a composite sample, keeping litter and soil layers separate. Soil mesofauna were extracted separately from litter and soil samples for five days using Tullgren extraction methods. Extracted animals were preserved in 75% ethanol and subsequently grouped taxonomically. Macrofauna were identified to the order or family level, while mesofauna were identified to the family or genus level, and allocated to trophic guilds. Because omnivory is common among soil invertebrates, most guilds were assigned to multiple food sources (live plant material, litter, bacteria, fungi, soil organic matter, and other invertebrates) based on current knowledge (Potapov *et al*. 2022). The body length and width of all individuals, or of at least 20 randomly selected individuals per guild, plot and layer when more were present, were measured to the nearest 0.1 mm using an ocular micrometer integrated into a dissecting microscope (Schwarz *et al*. 2017), and fresh body mass was estimated with group-specific allometric regressions based on body length and width (Hale *et al*. 2004; Mercer *et al*. 2001; Petersen 1975; Sohlström *et al*. 2018) (see Table S4 for formulas). Density was expressed per m^2^ by dividing abundance by the sampled area. Where all individuals were measured, guild biomass was calculated as the sum of individual body masses; otherwise, it was calculated as density multiplied by the mean body mass of the measured individuals. In total, 32 guilds of soil meso- and macrofauna were identified, which together with the five nematode trophic groups formed the 37 nodes of the reconstructed food webs (raw data are available from figshare; Data availability).

### Food-web reconstruction and energy flux estimation

We employed a ‘multichannel’ food-web reconstruction approach (Potapov 2022), integrating all trophic guilds of free-living soil nematodes, meso- and macrofauna into a unified dataset. For each guild, we used information from the reconstruction on feeding preferences (plants, algae, litter, fungi, bacteria, soil organic matter, or invertebrate predation), spatial niche (soil and litter), protection traits (traits of prey that reduce their likelihood of being preyed upon), and C and N content traits (proportion of carbon and nitrogen in the body) at supraspecific taxa level (Potapov 2022). We applied generic rules of food-web reconstruction based on food-web theory to infer weighted trophic interactions among all nodes, with the following theoretical rules (Potapov 2022): (1) feeding interactions between soil animal taxa are shaped by phylogenetically inherited variation in feeding preferences and predation abilities, as reflected by the resource preferences recorded in the trait table (Potapov *et al*. 2022); (2) predator-prey interactions are mainly governed by the optimal predator-prey mass ratio (PPMR) (Brose *et al*. 2019), in which predators are typically larger than their prey, although this ratio can be altered by specific characteristics such as hunting strategies or parasitic behaviour (Potapov 2022); (3) the strength of trophic interactions is influenced by the degree of spatial niche overlap, especially in terms of vertical distribution (Potapov 2022); (4) predators tend to favour more abundant prey due to higher encounter rates (Gauzens *et al*. 2019); (5) protective characteristics of prey, whether physical, chemical or behavioural, reduce the probability of predation (Peschel *et al*. 2006). By integrating these rules, we constructed the most plausible trophic interaction matrix, assigning feeding preferences and prey selection based on existing literature and considering factors such as size, spatial niche, total biomass, and prey protection traits. These data were then merged with our measurements of body mass and biomass per m^2^ to reconstruct the soil food web. Soil food webs were reconstructed separately for each plot (Figs. 2 and S5 to S10); plot values were then analysed with sites as independent replicates (Figs. 2 and S5 to S10). Nematodes were assigned to micro-food webs and all other soil animals, including mesofauna, to macro-food webs (Potapov *et al*. 2021).

To calculate energy fluxes within the food web, we utilized the *fluxweb* package (Gauzens *et al*. 2019), which incorporates reconstructed interaction networks, biomass, body mass, metabolic losses, and soil temperature. Metabolic rates per unit biomass were estimated from the mean fresh body mass of each guild and the mean soil temperature of each ecosystem during sampling, using group-specific metabolic regressions (Ehnes *et al*. 2011) (coefficients in Table S5). Diet-specific assimilation efficiencies were derived from the nitrogen content of each prey or basal resource, with different efficiencies depending on the resource type, which varies widely between consumer groups (Jochum *et al*. 2017; Potapov 2022). Energy fluxes to each node were then calculated, taking into account metabolism per biomass, assimilation efficiencies and predation losses, assuming a steady-state system where energy losses are compensated by lower trophic levels (Barnes *et al*. 2018; Gauzens *et al*. 2019). Finally, the fluxing function was applied to the reconstructed interaction networks to estimate the energy fluxes between all the nodes of the food web (Potapov *et al*. 2024), with the results being expressed in mW m^−2^.

### Soil food-web parameters, energy channels, and ecological indicators

We calculated bulk parameters and food web indicators for each plot to analyse changes in soil micro- and macro-food webs along secondary succession. The bulk parameters included: (1) total fresh biomass of all surveyed animal groups (g m^−2^); (2) total energy flux (mW m^−2^); (3) the ratio of total biomass to total energy flux (B/E), which quantifies the standing biomass maintained per unit energy throughput and thus approximates turnover time (Odum 1969); (4) scaled animal richness (given the different sampling methods for nematodes, meso- and macrofauna, we calculated total multitrophic taxonomic richness according to ref. (Allan *et al*. 2014), by averaging the standardised values for each group). Energy channels were classified by food source: herbivory (outgoing fluxes from living plants), algivory (from algae), litter and soil consumption (from plant litter and soil organic matter), bacterivory (from bacteria), fungivory (from fungi), and predation (from all animal nodes) (Potapov 2022). We further calculated four food web indicators: (1) the ratio of herbivory and algivory to litter and soil consumption, as a proxy for the balance between ‘green’ (herbivory-dominated) and ‘brown’ (detritivory-dominated) energy pathways (Wolkovich *et al*. 2014); (2) the ratio of predatory to basal resource consumption, as a proxy for top-down control (Rosenberg *et al*. 2023); (3) the ratio of bacterivory to fungivory, a proposed proxy for relatively fast versus slow energy and nutrient turnover (Rooney & McCann 2012); and (4) the ratio of faeces production (i.e., unassimilated food within the food web, calculated by multiplying all energy fluxes by the inverse of assimilation efficiency and summing the results) to soil organic matter consumption, serving as a proxy for carbon balance (Potapov *et al*. 2024). The last indicator is supported by three key points: soil invertebrates’ conversion of plant material to faeces boosts microbial biomass, microbial biomass production is crucial for soil organic matter formation and stabilisation, and invertebrates consuming soil organic matter, including associated microbial biomass, counteract the first two effects (Potapov *et al*. 2024). Total energy flux, the predation-to-basal consumption ratio and the bacterivory-to-fungivory ratio were calculated separately for micro- and macro-food webs.

### Statistical analyses

Statistical analyses were performed in R version 4.2.2 (R Development Core Team, 2022) with RStudio interface v. 1.4.1106 (RStudio, PBC).

To test Hypothesis 1 (H1), we employed a combination of linear mixed-effects models and partial least squares path modelling (PLS-PM) to examine changes in energy channels and their relationships with corresponding resources. We constructed linear mixed-effects models using the *lme4* package (Bates *et al*. 2015) to assess changes in bulk parameters and energy channels and soil food-web indicators along secondary succession, with ecosystem and river basin (as a block) as fixed effects and site (river basin × ecosystem; three plots per site) as a random effect (results presented in Table S6). Where necessary, variables were log10-transformed to meet the assumptions of normality. Using the *emmeans* package (Lenth 2025) with Kenward–Roger degrees of freedom (Kenward & Roger 1997), we tested linear and quadratic trends along the natural successional sequence (grassland, shrubland, secondary birch forest, primary forest), and compared the spruce and larch plantations with the secondary birch forest of the same successional stage. To account for multiple testing, P values were adjusted using the false discovery rate (Benjamini & Hochberg 1995). We used PLS-PM to include pathways testing both direct and indirect effects of basal resources, abiotic soil variables, and multitrophic animal diversity on energy flux. PLS-PM is a data analysis method that synthesises observed variables into latent constructs, assuming linear relationships among these constructs (Sánchez 2013); models were fitted with the *plspm* package. For instance, abiotic soil properties such as pH, soil temperature, and soil C/N ratio can be selected within the model to represent an overarching latent variable, encapsulating abiotic soil characteristics. The model excluded variables with loadings less than 0.7. Consequently, litter quality was represented by litter C/N ratio and cellulose concentration, and abiotic soil variables were represented by pH, soil temperature and soil C/N ratio. The succession stage was included as a continuous numerical variable in the path analysis, reflecting the ecological progression from early to late stages of ecosystem development (grassland = 1, shrubland = 2, secondary forests = 3, primary forest = 4). Because the three secondary forest types (birch, spruce and larch) developed over the same period, they were assigned the same value rather than an arbitrary sequential order. As a sensitivity analysis, the path model was refitted using only the natural successional sequence (grassland = 1, shrubland = 2, secondary birch forest = 3, primary forest = 4), excluding the two plantation forests (Table S8). The significance of path coefficients was assessed with t-tests, and their stability with 95% bootstrap confidence intervals (1,000 resamples). Goodness-of-fit (GOF) statistics were used to assess the predictive power of the model.

To test Hypothesis 2 (H2), we employed linear mixed-effects models to analyse total energy flux and two specific food web indicators, calculated separately for soil macro- and micro-food webs. The response variables included total energy flux, the predatory-to-basal resource consumption ratio, and the bacterivory-to-fungivory ratio (all log10 transformed). Models included ecosystem, food web (macro vs. micro), their interaction and river basin as fixed effects, and site, site × food web and plot as random effects. The ecosystem × food web interaction was tested with a Kenward–Roger F test to assess whether successional changes differed between food webs. The contrasts described above were tested within each food web, and an additional contrast tested whether the linear contrast differed between macro- and micro-food webs (Table S7).

To assess whether the results depended on key assumptions of the food-web reconstruction, we performed two sensitivity analyses, keeping all other reconstruction parameters constant. First, because metabolic rates, and thus energy fluxes, increase exponentially with temperature, we recalculated energy fluxes with the temperature differences among ecosystems halved and with all ecosystems set to the mean temperature (16.7 ℃), and retested the main results with the models and contrasts described above (Fig. S12). Second, to account for uncertainty in feeding preferences, we varied the weight of auxiliary food sources in the diets of trophic guilds (omnivory parameter; 0.2 in the main analysis) from 0 to 1 in increments of 0.1 and compared energy fluxes among ecosystems (Fig. S13). To identify which taxa drove the increase in B/E, we calculated for each taxon and ecosystem its share of total animal biomass and its energy flux per unit biomass (energy flux into the taxon divided by its biomass, both summed across plots). We then recalculated B/E for each plot after excluding earthworms (Lumbricina), using the energy fluxes of the reconstructed food webs, under both the baseline and common (16.74 °C) temperatures. B/E was tested with the models and contrasts described above (Fig. S14).

## Data and code availability statement

The data and code that support the findings of this study are openly available in Figshare at.

## Acknowledgements

This research was supported by the National Natural Science Foundation of China (Nos. 32572029; 32171756; 42677404), Science and Technology Projects of Xizang Autonomous Region, China (No. XZ202501JX0012) and Sichuan Science and Technology Program (Nos. 2026NSFSC0131; 2026NSFSC0975). AP was funded by the Deutsche Forschungsgemeinschaft (DFG, German Research Foundation) – Project number 493345801. NE acknowledges support from the DFG (FZT 118, 202548816; Ei 862/29-1; Ei 862/31-1). AZ was funded by the China Scholarship Council (No. 202304910477).

## AUTHOR CONTRIBUTIONS

X.P., A.M.P. and A.Z. conceptualized the idea and designed the study. A.Z., C.T., Y.W., R.L. and Z.C. collected data. A.Z. compiled the data, did data analysis and wrote the manuscript with supervision from A.M.P. and X.P. N.E. and K. H. contributed to the interpretation of the results and edited manuscript drafts. J.L., Z.Z. and B.Z. contributed to statistical analysis and edited manuscript drafts. All authors have read, reviewed and agreed to this version of the manuscript.

